# Short Excitatory Projections Promote Burst Activity and Modular Functional Organization in Neural Networks

**DOI:** 10.64898/2025.12.11.693643

**Authors:** M. Sharif Hussainyar, Di Yun, Dong Li, Ji-Song Guan, Claus C. Hilgetag

## Abstract

Physical constraints in the brain favor short, distance-dependent connections. However, short-range projections are not merely a compromise imposed by these constraints; they also serve essential functional roles. Here, we explore networks with balanced excitation and inhibition, demonstrating that short excitatory projections fundamentally increase the number of common presynaptic inputs, thereby enhancing the correlations of neural responses. Strengthened recurrent correlations can lead to bursting states, amplifying postsynaptic neuronal activity and supporting signal propagation over long distances within the network. Moreover, even without an explicitly imposed modular structure, short excitatory connections can give rise to functional clustering and support a broad spectrum of modular functions. Therefore, this structural principle provides a mechanistic basis for functional column formation and dynamic burst propagation, suggesting that the brain transforms structural wiring constraints into opportunities for complex computation and distributed processing. These findings pave the way for a deeper understanding of how the brain transforms structural limitations into opportunities for complex computation.

## 1. Introduction

Neural circuits of the cerebral cortex are organized in intricate patterns of connectivity that span multiple spatial and functional scales. Across all levels, neural connectivity is neither purely regular nor entirely random, but instead exhibits complex patterns of organization. Previous studies have identified structural features, such as small-world topology [1, 2], scale-free degree distributions [3], and modularity [4, 5], etc., that are often associated with functional advantages in information processing [6–9]. Among these, modular organization has served as a particularly influential framework for understanding cortical networks [10, 11].

Although modularity may result from mechanisms such as synaptic plasticity [12, 13] or activity-dependent refinement, such outcomes still rely on relatively intricate processes. This raises a fundamental question: do cortical circuits explicitly need to possess a modular structural organization to gain the functional benefits of modularity, or can simpler organizational principles achieve the same benefits?

Among the simplest yet highly influential principles shaping network architecture is the distance over which neural connections are formed. Cerebral networks comprise both long-range and short-range projections. While long-range projections have been extensively studied for their role in inter-regional communication and functional integration [14–16], short-range connections remain less well understood. This is particularly important because cortical circuits show a clear pattern of connection probability decreasing with distance, creating dense local networks that vastly outnumber sparse long-range projections [17–21].

Direct functional measurements using simultaneous whole-cell recordings reveal substantial local excitatory connectivity [22]. Further studies showed that connections from excitatory neurons are spatially constrained, with synapses typically located at a mean distance of only 67 *µ*m from the postsynaptic soma and predominantly targeting basal dendrites [23]. Similarly, connection probability in rat layer 2/3 neocortex decreases systematically from 0.09 within ±25 *µ*m to 0.01 at distances >100 *µ*m [24], establishing that Short Excitatory Projections (SEP) represent a fundamental organizational principle of cortical circuits.

Crucially, this type of connectivity is not merely a structural compromise imposed by spatial constraints. Many excitatory cell types, including layer 4 spiny stellate cells and near-projecting neurons, have characteristically short axonal arbors and operate primarily within local columns or areas [20]. These neurons differ from long-projecting counterparts in their distinct gene expression profiles, connectivity patterns, firing properties, and developmental trajectories [24], indicating their functional specialization, rather than developmental truncation.

Accordingly, theoretical frameworks for spatially extended neural networks have established that spatial connectivity patterns can fundamentally alter network dynamics [25–27]. For example, when inhibitory connections span broader distances than excitatory ones, networks can undergo symmetry-breaking bifurcations that enable complex computations [28]. However, the complementary question, how constraining excitatory connections to short projections affects network dynamics, remains underexplored. Given that cortical circuits exhibit predominantly SEPs [20,22,24], understanding the functional implications of short-range constraint is essential for developing a coherent theoretical framework of cortical computation.

The prevalence of distance-dependent connections, specifically SEPs, also relates to the question of whether simple connectivity principles can provide functional benefits of modularity, in the absence of intricate structural modularity. This question is particularly relevant for the fundamental feature of cortical columns, where functional columnar organization may emerge from distance-dependent connectivity principles rather than actually requiring discrete anatomical modules.

### 1.1. Local Connectivity as a Structural Basis for Functional Organization

The existence and functional relevance of cortical columns have long been debated. Some classic studies argue that anatomical microcolumns provide the structural basis for functional specialization [29, 30], while others suggest that functional organization arises from experience-dependent network dynamics rather than discrete anatomical units [31, 32]. In contrast, other perspectives question the functional role of columns altogether, emphasizing that cortical processing is better understood as emerging from distributed, flexible ensembles rather than rigid modules [33–36].

Despite these debates, there is broad agreement that functional specificity exists across cortical areas, even if its structural substrate remains uncertain. Increasing evidence suggests that this specificity may arise not from fixed microcolumns, but from the arrangement of local connectivity, particularly SEPs. Understanding how this type of connectivity might give rise to functional specificity, even without explicit anatomical modularity, may thus reconcile the long-standing columnar debate and illuminate fundamental principles of cortical circuit design.

### 1.2. Burst Dynamics Arising from Local Recurrent Connectivity

Local connectivity not only supports columnar organization but also fundamentally shapes burst dynamics. In biological networks, the density and spatial organization of recurrent excitatory connections determine whether local circuits generate localized, self-terminating bursts or sustained propagating activity. For example, hippocampal CA3 regions with high recurrent connectivity generate robust bursts, whereas seizures originate from the CA1 region with fewer recurrent connections [37].

Recent mesoscale studies of cultured cortical networks have revealed that spontaneous burst activity originates from specific zones characterized by enhanced recurrent connectivity and structural transitions [38]. Critically, these burst initiation zones emerge at network boundaries and cluster edges where anisotropic connectivity patterns create local recurrent loops, rather than in regions of uniform high density. This spatial specificity suggests that the interplay between local recurrent circuit architecture and network-wide connectivity patterns, rather than absolute connectivity density alone, determines where and how bursts arise.

Despite these observations, the circuit-level mechanisms linking local connectivity structure to burst generation and propagation remain unclear.

Addressing these questions connects directly to the debate on the organization of cortical columns above. Functional columns may emerge from local connectivity patterns rather than discrete anatomical modules. Similarly, burst dynamics may arise from fundamental principles of recurrent circuit organization.

Here, we hypothesize that short excitatory projections function as correlation amplifiers. They may enhance signal propagation via two mechanisms: (1) increasing shared presynaptic inputs among neighboring neurons, which elevates response correlations, and (2) promoting burst-like activity in local recurrent circuits that amplifies weak signals for long-range transmission.

Our computational modeling systematically varies SEP density to examine how these connections shape recurrent amplification, burst generation, and network-wide propagation. The results demonstrate that SEPs are not redundant wiring elements but represent essential components for efficient network communication by enhancing both local activity representation and global signal propagation.

These findings suggest a unified framework: even without explicit structural modularity, SEPs can equip cortical networks with the functional advantages of modular architectures by linking local connectivity, emergent dynamics, and cortical computation.

## 2. Methods

In this study, we combine two main components: (i) simulating neuronal membrane potential dynamics using a conductance-based Integrate-and-Fire model [39], and (ii) constructing neuronal networks with distance-dependent connections to capture spatial characteristics observed in biological circuits.

### 2.1. Modeling Neuronal Membrane Potentials

To simulate membrane potential dynamics, we employed the conductance-based Integrate-and-Fire model introduced by Vogels and Abbott [39]. The membrane potential dynamics of each neuron are governed by:

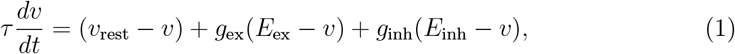

where *τ* is the membrane time constant, *v* is the membrane potential, *v*_rest_ is the resting membrane potential, *g*_ex_ and *g*_inh_ are the conductances of excitatory and inhibitory synapses, and *E*_ex_ and *E*_inh_ are the reversal potentials for excitatory and inhibitory synapses, respectively.

The synaptic conductances *g*_ex_ and *g*_inh_ evolve according to:

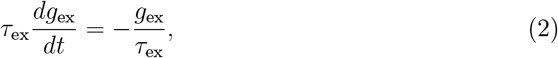

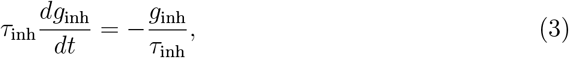

where *τ*_ex_ = 15.0 ms and *τ*_inh_ = 5.0 ms are the time constants for excitatory and inhibitory synaptic conductances, respectively. A spike updates the conductance by *δg* unless the postsynaptic neuron is in the refractory period (*p*_*r*_ = 2.5 ms), where the parameters *δg* are fixed as 0.2 (excitatory to excitatory), 0.45 (excitatory to inhibitory), 3.05 (inhibitory to excitatory), and 3.0 (inhibitory to inhibitory cells). To initiate and maintain network activity, random inputs with intensity *δg* = 0.2 are injected to a small subset of excitatory neurons at each time step.

### 2.2. Network Configuration and Connectivity

We simulated 5000 neurons randomly distributed within a two-dimensional (2D) space with side length *L* and periodic boundary conditions, where 80% of the neurons were excitatory and 20% inhibitory, mirroring the typical composition observed in biological neural circuits [40–43].

As a baseline configuration, we first implemented a network with purely random connections, independent of spatial distances between neurons. In this configuration, both excitatory and inhibitory neurons formed connections randomly with other neurons, producing an unstructured network with a uniform global connection probability of *p* = 0.1.

To model SEP, we randomly selected a proportion of the excitatory neurons (referred to as SEP neurons hereinafter) and rewired their outward connections based on the physical distance between pre- and postsynaptic neurons. The connection probability *p*_0_ between two neurons is given by:

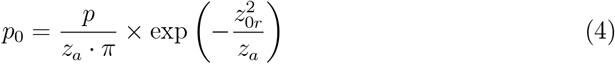

where *z*_*a*_ = 0.128*L* is a scaling parameter, *z*_0*r*_ is the distance between the two neurons, and *p* = 0.1 is the global connection probability. The number of outward connections was kept exactly the same as those of the baseline random network.

This modeling setup ensures that a specified proportion of neurons exhibit distance-dependent connectivity (SEP neurons), as illustrated in Figure 1A, while the remaining neurons (Non-SEP) form connections with distance-independent connectivity (Figure 1B). Importantly, the total number of outgoing connections per neuron is preserved across both SEP and Non-SEP populations, ensuring that differences in network dynamics arise solely from the spatial distribution of connections rather than from variations in output degree.

**Fig 1.**
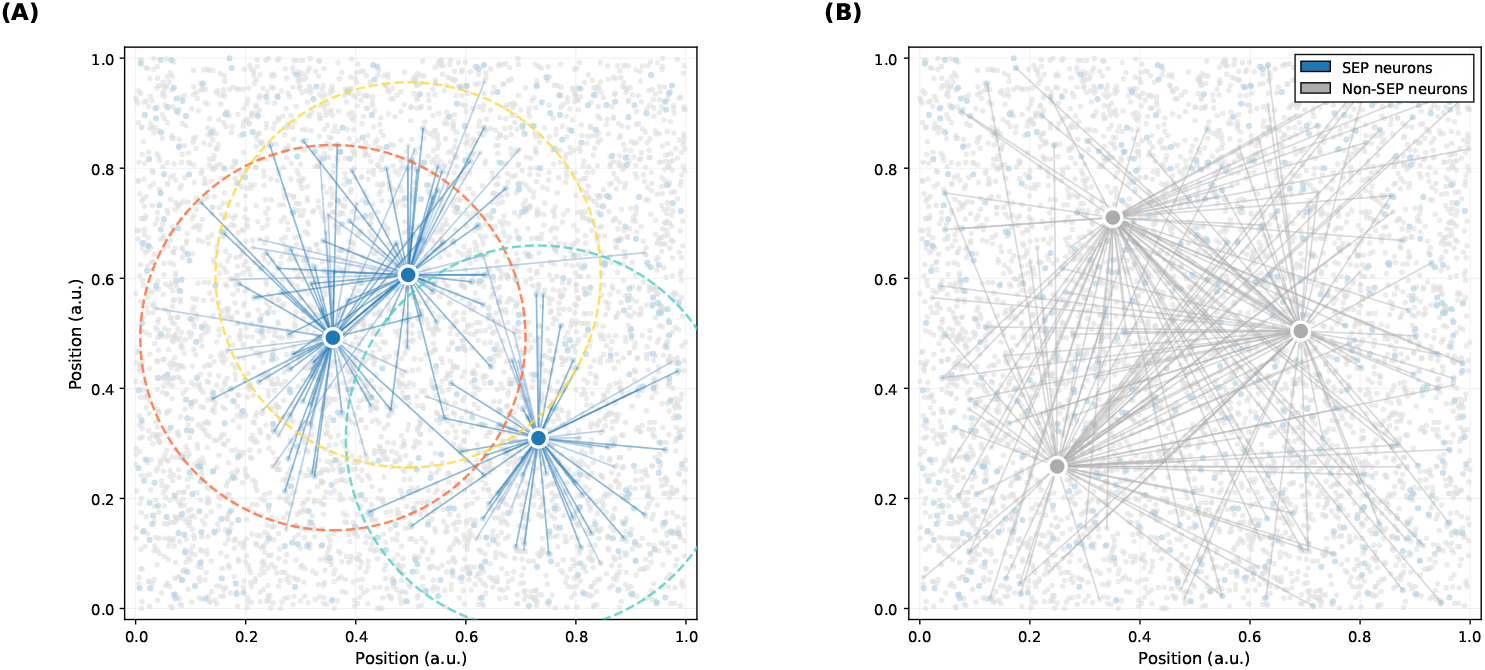
Connection Principles in the neural network model. The network comprises two distinct neuronal populations with different connection probability rules. (**A**) SEP neurons (blue) exhibit distance-dependent connection probability, preferentially forming synaptic connections with spatially proximate neighbors (dashed circles indicate regions of elevated connection probability). (**B**) Non-SEP neurons (gray) exhibit spatially uniform connection probability, forming connections with equal probability regardless of spatial distance. Highlighted neurons (larger, darker markers) represent example neurons whose outgoing connections are visualized.

We systematically studied the effects of different amounts of SEP neurons by gradually changing their number. As examples, we primarily present the cases of 0, 500, and 1000 SEP neurons (denoted as SEP=0, SEP=500, and SEP=1000, respectively).

### 2.3. Data Analysis

To assess how short-projecting connections modulate network responsiveness, we introduced targeted perturbations by stimulating a randomly selected neuron in networks with different numbers of SEP neurons. In each configuration, we stimulated the same neuron with identical intensity and duration to ensure consistent experimental conditions and studied its effect on every other neuron using the same procedure widely employed in experiments [44, 45]. For each neuron, the change in activity was quantified as: ΔFiring Rate = stimulated firing rate − baseline firing rate.

To investigate the underlying mechanisms of network responsiveness, we examined the temporal structure of individual neuronal firing patterns, focusing particularly on burst activity. Bursts were defined as sequences of at least three spikes with inter-spike intervals ≤ 7.5 ms, as these events are known to enhance synaptic transmission, increase signal reliability, and contribute to synaptic plasticity and learning. Additionally, functional connectivity was calculated based on correlations in the firing rates of neurons, and network clustering was identified using a consensus model for modularity maximization [46].

## 3. Results

Using the network configurations and analysis methods described above, we systematically examined how SEP density modulates network dynamics and information processing. In such an excitation-inhibition (E-I) balanced network, we observed that functional dynamics gradually evolve as the number of SEP neurons increases from a baseline of purely random connectivity. These dynamics include network responsiveness to external stimuli, signal propagation efficiency, burst activity patterns, functional connectivity, and overall firing rate characteristics.

### 3.1. Network Responsiveness and Signal Propagation

Increasing the number of SEP neurons significantly enhances the network’s responses to external stimuli and signal propagation on the network. Examples are shown in Figure 2, where networks with a higher number of SEP neurons (SEP = 1000) exhibited significantly greater changes in firing rate following perturbation compared to networks with fewer such connections (SEP = 500). Figure 2A reveals a clear linear correlation between baseline activity levels and response magnitude, with neurons in the SEP = 1000 network demonstrating both higher baseline activity and larger perturbation-induced changes. Panel Figure 2B further illustrates the broader dynamic range in the SEP = 1000 network through probability distributions of firing rate changes (plotted on a logarithmic scale).

**Fig 2.**
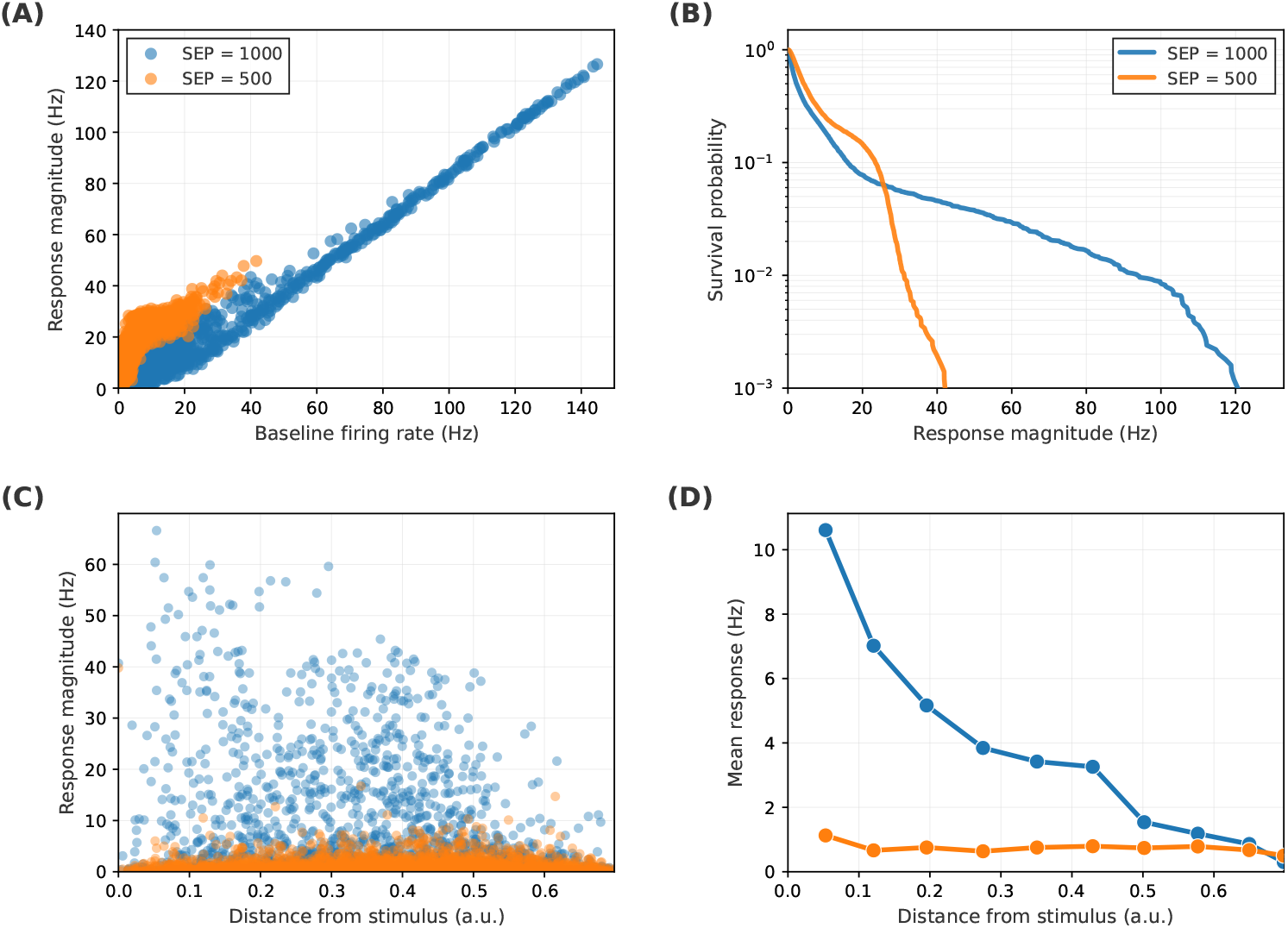
Neuronal responses to single-neuron perturbations: comparing networks with different densities of short excitatory projections (SEP). **(A)** Baseline firing rates versus perturbation-evoked response magnitudes, showing that networks with more short-range excitatory connections (SEP = 1000, blue) exhibit larger responses than networks with fewer such connections (SEP = 500, orange). **(B)** Survival probability *P* (response ≥ *x*) of perturbation-evoked response magnitudes, indicating a broader high-response tail in the high-SEP network. **(C)** Response magnitude as a function of distance from the stimulated neuron, showing enhanced signal propagation in SEP = 1000 networks. **(D)** Distance-binned mean response magnitude, demonstrating that higher-SEP networks maintain elevated responses over greater distances from the stimulus.

The signal propagation follows the same pattern. Panels C and D in Figure 2 demonstrate how the effects of localized perturbations spread across the network as a function of distance from the stimulated neuron, and it is clear that networks with a higher number of SEP neurons (SEP = 1000) exhibit significantly enhanced signal propagation compared to those with fewer SEP (SEP = 500). In the SEP = 1000 network, substantial changes in firing rates are observed even in neurons at greater distances from the perturbed neuron, indicating more widespread propagation of the perturbation effect. By contrast, in the SEP = 500 network, the impact remains more localized, with firing rate changes diminishing rapidly with distance.

### 3.2. Burst Activity and Emergence of Functional Clusters

The mechanisms underlying enhanced responsiveness and signal propagation in networks with short excitatory projections are fundamentally shaped by their bursting dynamics. Networks with more SEP neurons (SEP = 1000) generate significantly more frequent and longer bursts than those with fewer such connections (SEP = 500). Figure 3 illustrates these findings across four panels. Panels A and B display raster plots of selected excitatory (red) and inhibitory (blue) neurons for networks with SEP = 1000 and SEP = 500, respectively. Panel C presents the distribution of burst counts per neuron (on a logarithmic y-axis), showing that networks with denser SEPs exhibit a pronounced rightward shift. Panel D shows the corresponding burst duration distributions. Neurons in the SEP = 1000 network not only burst more frequently but also sustain their bursts for longer periods, consistent with increased excitatory drive and greater local input correlation.

**Fig 3.**
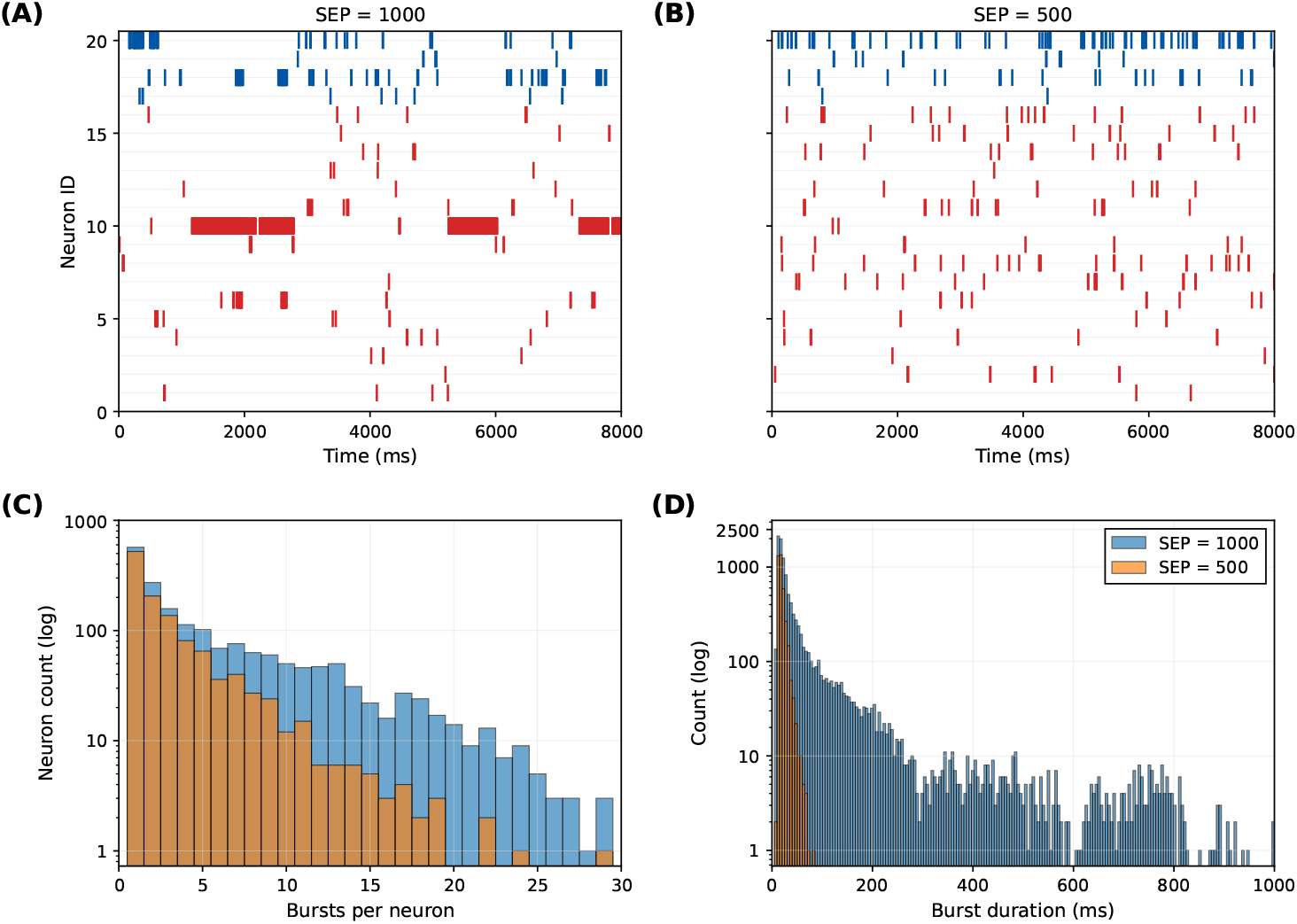
Burst activity analysis in neural networks with different short excitatory projections (SEP). **(A)** Spiking patterns of selected excitatory (red) and inhibitory (blue) neurons in networks with SEP = 1000, showing prominent burst activity. **(B)** Corresponding spiking patterns in networks with SEP = 500, displaying less synchronized firing. **(C)** Burst count distributions across neurons (log scale), demonstrating that networks with higher short excitatory projections generate significantly more bursts per neuron. **(D)** Burst duration distributions (log scale), showing that networks with SEP = 1000 not only produce more frequent bursts but also sustain longer burst episodes compared to networks with SEP = 500.

Figure 4 illustrates the spatial organization of burst activity patterns and their correlation with neuronal positioning in 2D space. Importantly, our network configuration as described in the Methods section contains no predefined structural clusters, and as evident in panels A and B of Figure 4, SEP neurons are uniformly distributed across the entire 2D space. Despite this homogeneous spatial arrangement of network architecture, the burst events exhibit distinct spatial clustering patterns that emerge spontaneously from the functional dynamics. These emergent functional clusters strongly correlate with the amount of SEPs, demonstrating that even uniformly distributed SEP neurons can generate spatially organized functional activity.

**Fig 4.**
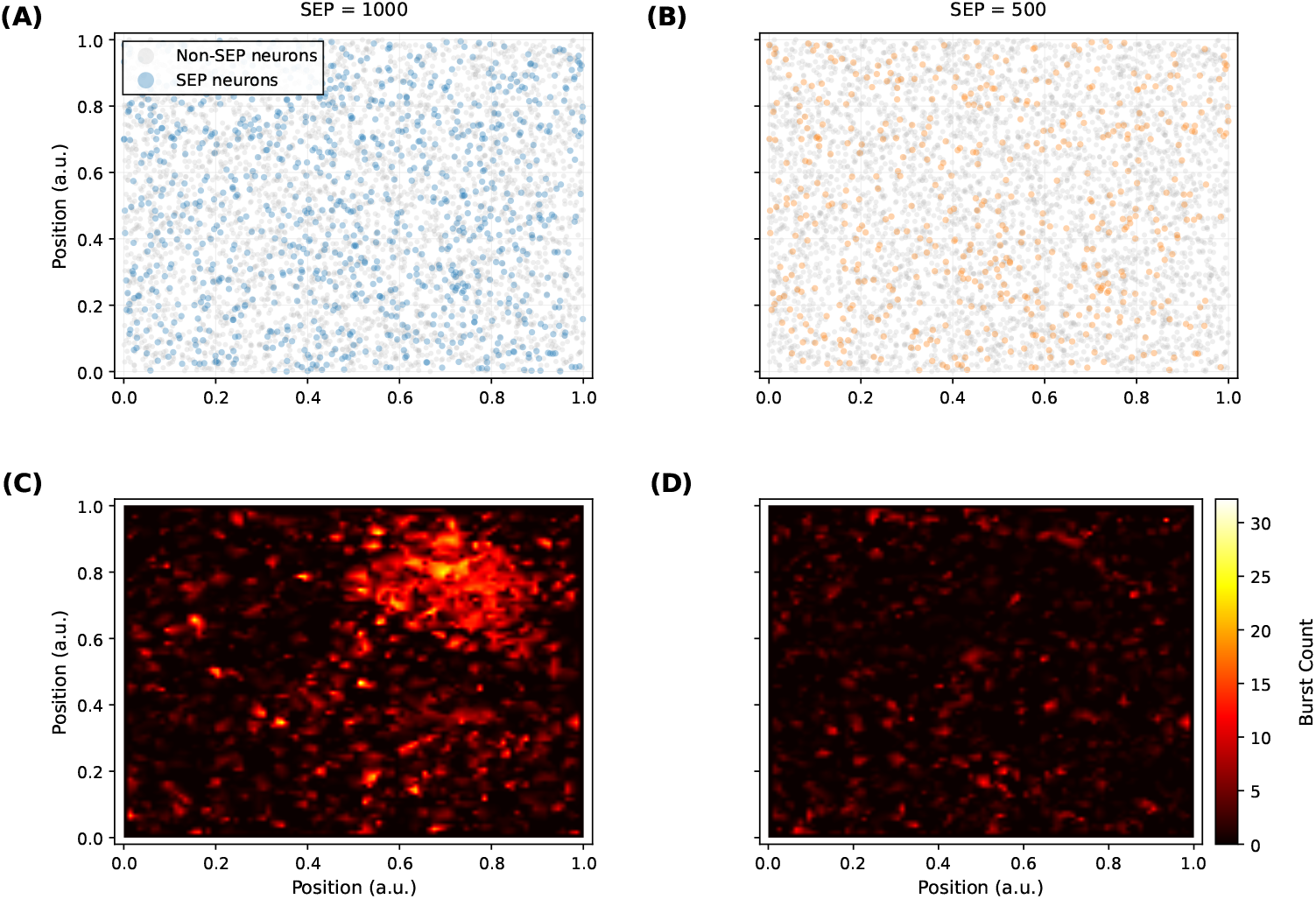
Spatial distribution of burst activity in neural networks with different short excitatory projections (SEP) configurations. **(A)** Spatial positions of SEP neurons (red dots, n=1000) and global neurons (light blue dots, n=4000) in the SEP = 1000 network, illustrating the distribution of short-projecting connections within the neural population. **(B)** Spatial positions of SEP neurons (red dots, n=500) and global neurons (light blue dots, n=4500) in the SEP = 500 network, showing the reduced density of short-projecting connections. **(C)** Burst count heatmap for networks with SEP = 1000, showing the spatial distribution of bursting activity across the neural field. Higher burst counts (yellow regions) indicate areas of intense synchronized activity. **(D)** Burst count heatmap for networks with SEP = 500, displaying reduced overall burst activity and more dispersed spatial patterns compared to the SEP = 1000 condition. The color bar indicates burst count values, with warmer colors representing higher bursting activity.

In fact, not only do burst events exhibit spatial clustering, but firing rate changes also display clustered spatial patterns (see Figure S1). Firing rate changes, burst frequency, and burst duration parameters are in fact strongly correlated (see Figure S2).

Obviously, the enhanced temporal dynamics observed in high-SEP networks arise not from anatomical clustering, but from the emergent self-organization of burst activity through local connectivity patterns. SEP facilitates recurrent, temporally clustered activity patterns through enhanced local input overlap and spatial clustering, so as to promote conditions conducive to bursting, which in turn appears to support more robust signal propagation and increased network responsiveness. The emerged dynamics are further evidenced by the functional connectivity (as two examples shown in Figure 5), where networks with SEP = 1000 exhibited stronger and more distinct functional clusters than those with SEP = 500, even though neither of the underlying structural connectivity is explicitly designed to be modular at all.

**Fig 5.**
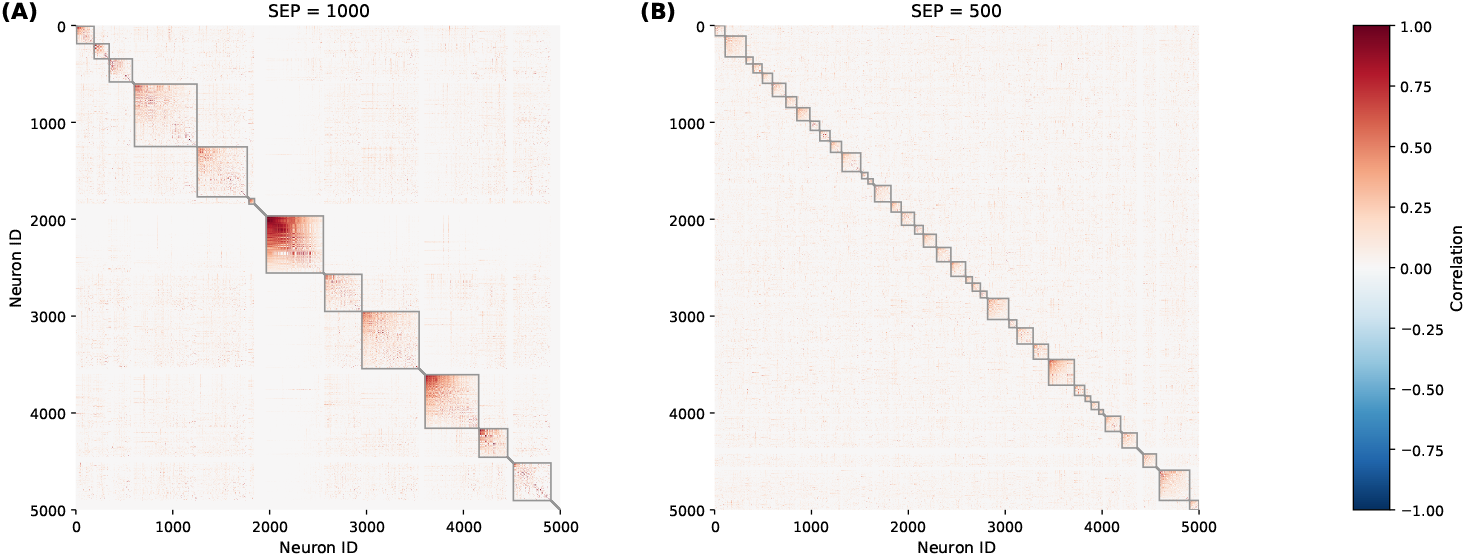
Functional connectivity of neuronal networks with varying numbers of short excitatory projections (SEP). **(A)** Network with SEP = 1000 showing larger and more distinct clusters of synchronized neuronal activity, indicating stronger functional synchronization. **(B)** Network with SEP = 500 displaying sparser connectivity patterns. Clusters emerge from the underlying activity patterns despite the absence of explicit structural modularity in the network architecture. The color bar indicates correlation values ranging from − 1 (blue, negative correlations) to 1 (red, positive correlations). Clusters were detected using the consensus model for modularity maximization [46].

### 3.3. Increased Network Excitability and Variability

The burst activity and functional clustering introduced by SEP are intrinsically linked to the altered overall excitability of the network. Figure 6 demonstrates how the average firing rate progressively increased with the number of neurons possessing short-projecting connections, rising from below 5 Hz to approximately 30 Hz (Panel A). This trend underscores the critical role of local connectivity in modulating network excitability.

**Fig 6.**
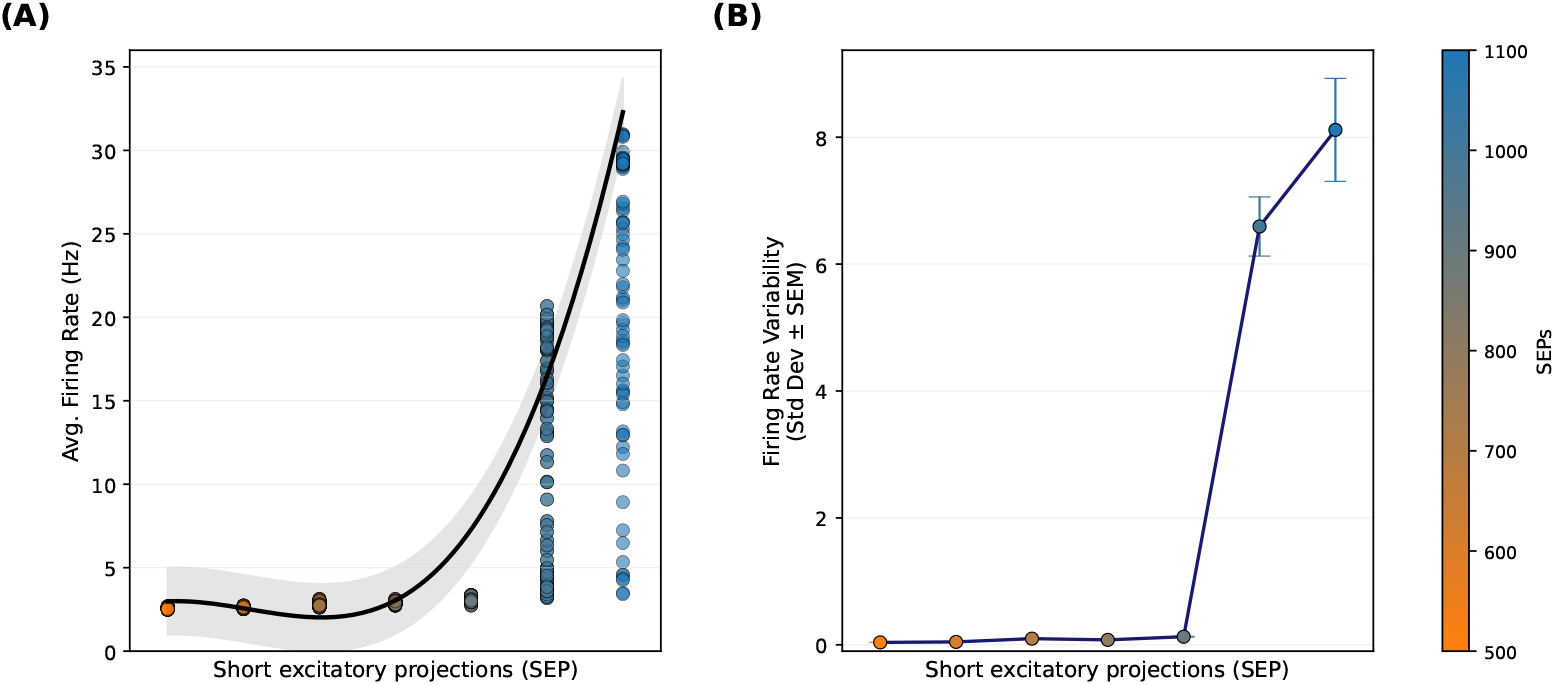
Firing dynamics across networks with increasing short excitatory projections (SEP). Each network was simulated over 100 trials with different initial conditions and is ordered on the x-axis by increasing SEP. **(A)** Firing rates of all neurons per trial and network are shown as scatter points. As the number of short excitatory projections (SEP) increases, the maximal firing rate per network also increases. A cubic polynomial fit to the maximal firing rate (solid line) with a shaded 95% confidence band illustrates this upward trend. **(B)** Trial-to-trial variability in firing rate for each network, quantified as the standard deviation of firing rates and plotted as mean ± SEM across trials. Variability increases with SEP, indicating greater heterogeneity in population activity at higher short excitatory connectivity.

Importantly, increased SEP also introduced greater trial-to-trial variability, as depicted in Figure 6B. The standard deviation of firing rates across trials increased with higher SEP levels, suggesting that the networks became more sensitive to initial conditions and local fluctuations. This enhanced variability reflects richer population dynamics.

The cubic polynomial fit to the firing rate data (Figure 6A) reinforces the non-linear relationship between SEP and network excitability. This non-linearity suggests the existence of critical thresholds in connectivity that, when crossed, lead to qualitative changes in network behavior. Such transitions are characteristic of complex systems and eventually lead to the emergence of the burst activity and functional clustering observed in our previous analyses.

### 3.4. Structural Mechanisms Underlying Enhanced Network Dynamics

The mechanism underlying this increased excitability appears to be related to the enhanced recurrent excitation facilitated by SEP. SEP neurons enable other neurons in the neighborhood to receive a higher proportion of common presynaptic inputs (Figure 7), creating a positive feedback loop that amplifies activity. Importantly, this local correlation amplification via shared inputs aligns with theoretical predictions that distance-dependent recurrent connectivity in balanced E–I networks generates spatially structured correlations [47], and thus provides a principled explanation for our observations. This local amplification, together with the network’s inherent excitatory–inhibitory balance, creates conditions favorable for both higher baseline activity and a greater dynamic range in response to perturbations.

**Fig 7.**
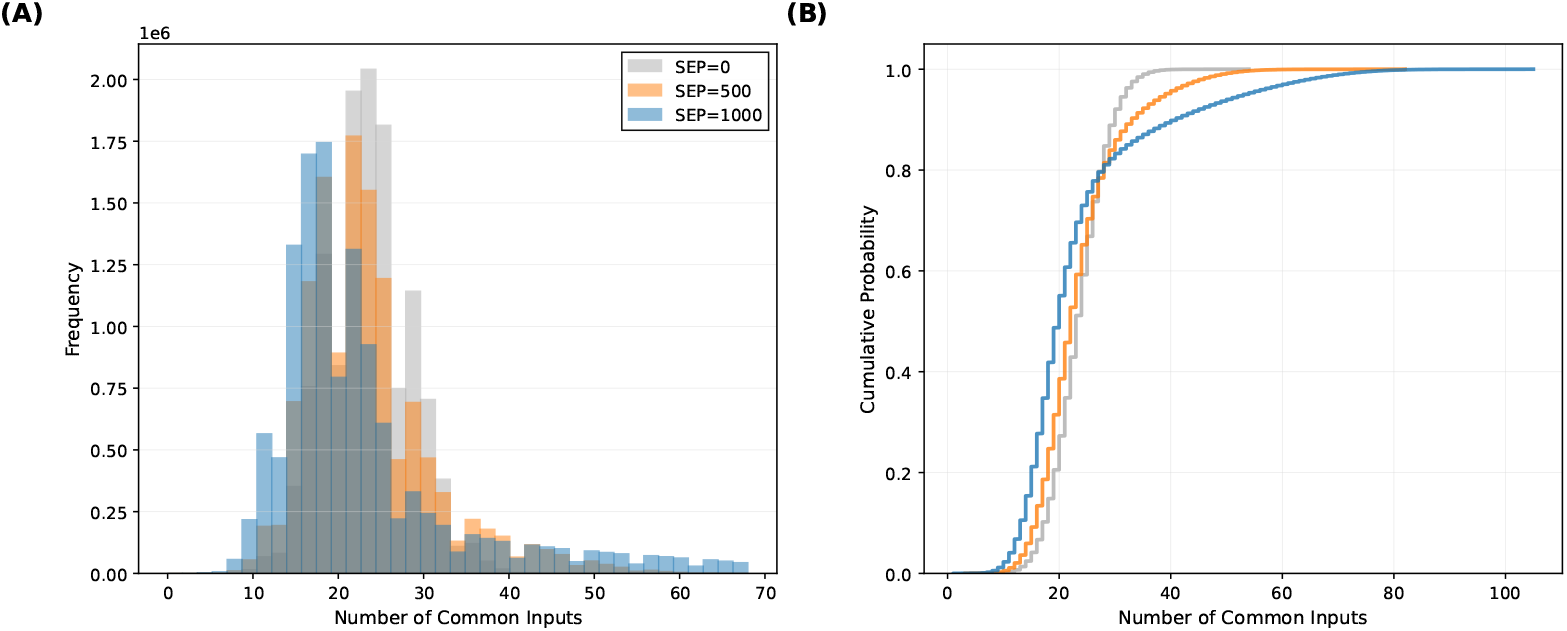
Short excitatory projections increase shared presynaptic input among neuron pairs. **(A)** Distributions of common excitatory presynaptic inputs for all neuron pairs with at least one shared input in networks with 0, 500, or 1000 SEP neurons. Increasing the number of SEP neurons systematically shifts the distribution toward larger numbers of common inputs. **(B)** Cumulative distribution functions of common excitatory presynaptic inputs for the same networks, demonstrating a progressive rightward shift with higher SEP, consistent with enhanced local input overlap and recurrent excitation.

## 4. Discussion

The present study highlights the fundamental importance of SEPs in neural networks, demonstrating that these connections are far from being merely structural redundancies or byproducts of physical constraints. Instead, our findings reveal that SEPs serve as critical architectural elements that significantly enhance network dynamics, signal propagation, and computational capabilities.

In balanced networks, incorporating SEPs fundamentally altered network behavior by increasing the number of shared presynaptic inputs among postsynaptic neurons. This structural modification enhanced correlated activity across postsynaptic populations, aligning with existing findings on spatial clustering and neural response synchronization [21]. The resulting functional network exhibited increased clustering coefficients and path length, indicating heightened local connectivity while maintaining a predominantly random network structure. Similar mechanisms have been reported in spatially extended excitatory–inhibitory spiking networks, where narrower-range excitatory connections compared with broader inhibitory projections, together with neural heterogeneity, disrupt coherent spatiotemporal patterns and give rise to reliable, input-slaved transient dynamics for robust computation [48]. In line with this work, our results show that introducing short excitatory projections into an otherwise largely random balanced network similarly enriches the dynamics and enhances functional clustering and signal propagation, without imposing an explicitly modular anatomical architecture. This balance demonstrates that SEPs can introduce localized clustering of activities without necessitating a modular or small-world topology.

Our findings offer a mechanistic resolution to the debate on the existence of anatomical cortical columns, as functional columnar organization can emerge directly from SEP patterns without requiring discrete anatomical microcolumns. In visual cortex, such local connectivity supports orientation tuning [49], and in somatosensory cortex it enables coordinated activity across whisker maps [50]. These findings demonstrate that distance-dependent connectivity principles generate functionally essential features; yet many theoretical models still treat local connectivity as a byproduct of spatial wiring constraints [51].

Our simulations show that spatial clustering of burst activity, clustered functional connectivity (Figure 4 and Figure 5), and enhanced response correlations (Figure S2) observed in our networks, despite the absence of predefined structural modules, demonstrate that SEPs create functional columns through a simple mechanism: by systematically increasing common presynaptic inputs among neighboring neurons (Figure 7). This enrichment generates local ensembles that mirror the coordinated activity characteristic of columnar processing in sensory cortices. Importantly, while SEPs do increase structural modularity (Figure S3), the resulting modularity values remain substantially lower than explicitly designed modular networks, indicating that functional columnar organization emerges through dynamic activity patterns rather than strict anatomical modularity.

In addition, our results align with experimental findings from cultured cortical networks [38], which demonstrated burst initiation and propagation at the mesoscale. Our results provide a mechanistic explanation for these mesoscale observations: at the local circuit level, SEP systematically increases common presynaptic inputs (Figure 3), creating the recurrent amplification circuits that manifest as burst initiation zones at larger scales and propagate over longer distances (Figure 2). This spatial clustering of burst activity can be attributed to recurrent signals combined with common input patterns, where neurons sharing similar inputs tend to activate together through recurrent connectivity. This mechanism paves the way to understand several phenomena observed in biological neural networks; for example, they exhibit robust tolerance, enabling meaningful responses even when cells are not individually targeted, as long as activation occurs within a defined range.

From an evolutionary perspective, our findings support the view that neural circuits have been optimized over time to meet both structural and functional demands. The optimization process involves minimization of conduction delays, passive cable attenuation, and wire length while maximizing synaptic density [51]. This evolutionary optimization suggests that SEPs are integral to efficient circuit functioning and represent a fundamental design principle rather than a mere byproduct of physical constraints. By showing that basic organizational principles alone can give rise to similar advantages, our work points toward a broader and more parsimonious framework for understanding how functional capabilities may emerge in the brain.

In conclusion, our findings contribute new insights into the role of SEPs in neural networks, supporting their evolutionary significance and functional impact. The results emphasize that SEPs, although operating within physical constraints, are essential to the dynamic responsiveness, computational complexity, and adaptive capabilities of neural networks. By increasing common presynaptic inputs and enhancing local recurrent connectivity, SEPs may provide a circuit-level explanation for both the emergence of functional columnar organization and the spatial patterns of burst initiation and propagation observed in cortical circuits. Future research could extend this framework to incorporate additional biological factors such as plasticity mechanisms, neuromodulation, and developmental constraints, further elucidating how short-projecting connections support cognitive processing, learning, and stability in neural systems. Understanding these mechanisms will be crucial for advancing both theoretical neuroscience and the development of biologically-inspired artificial neural networks.

## Acknowledgements

This work is supported in part by the following grants: LFF-FV76 to MSH, NSFC (32400853) to DY, NSFC(62061136001)/DFG SFB TRR-169 - A2 to JSG and CCH, NSFC(32225023) to JSG, and SFB 936-178316478-A1, SFB 1461/A4, and SPP 2041/HI 1286/7-1 to CCH. The funders had no role in study design, data collection and analysis, decision to publish, or preparation of the manuscript. We thank Arnaud Messé and Mariia Popova for helpful comments on the manuscript.

## Supplementary Materials

### Spatial structure of perturbation-evoked responses

Figure S1 illustrates the spatial distribution of perturbation-evoked firing rate changes for three example neurons located at different positions in the 2D network. For each neuron, we compared networks with high versus low short excitatory projections (SEP = 1000 vs. SEP = 500). After a single-neuron perturbation, the change in firing rate was computed for every neuron by subtracting the baseline firing rate from the stimulated condition and mapping these values back into physical space.

**S1.**
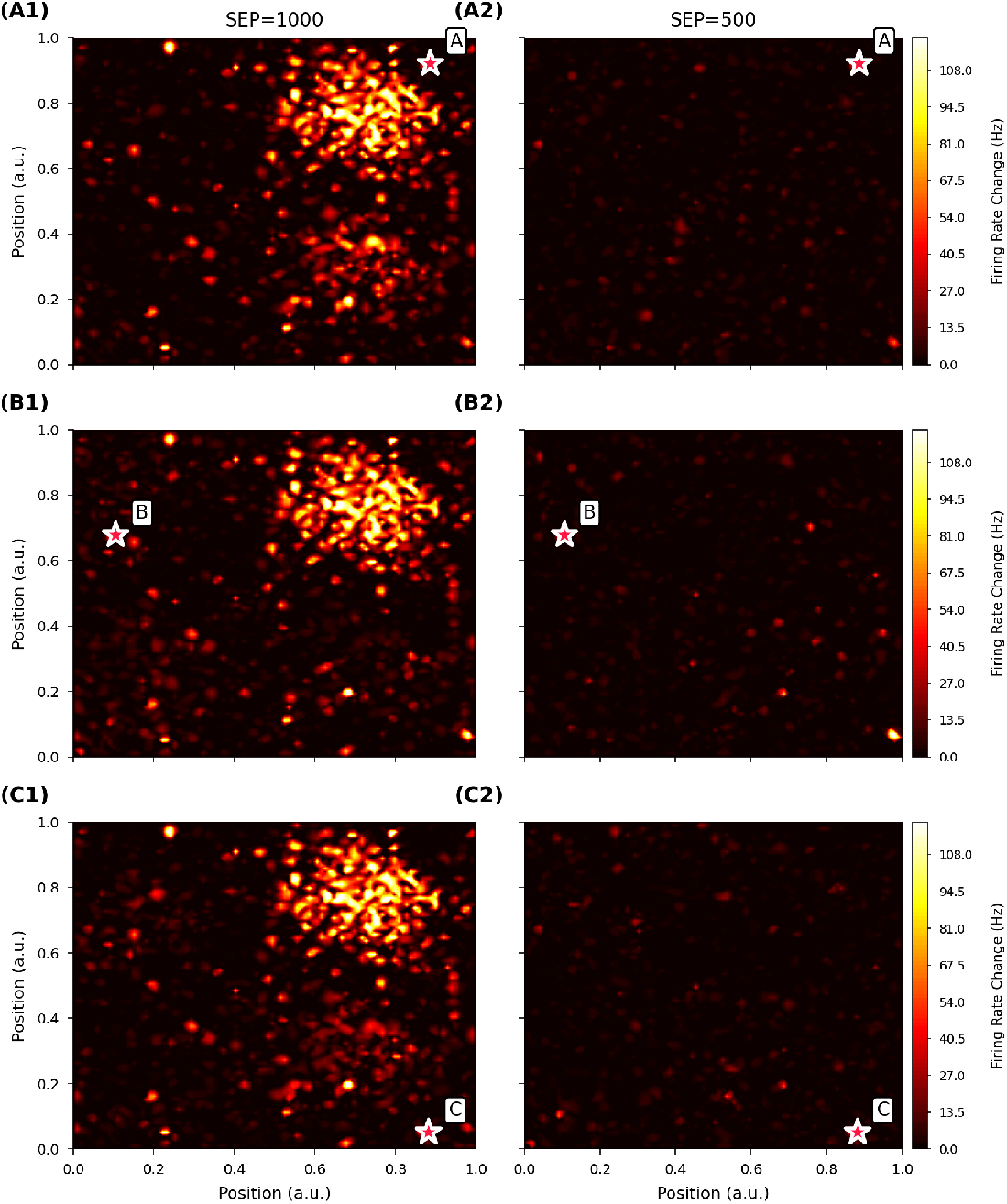
Spatial distribution of firing rate changes following single-neuron perturbations. Heat maps show the absolute change in firing rate (Hz) between baseline and stimulated conditions for three different perturbed neurons (rows A–C) in networks with high and low short excitatory projections (left: SEP = 1000; right: SEP = 500). The perturbed neuron in each panel is marked by a black star. Under SEP = 1000, perturbations generate extended regions of elevated firing rate change, with clear spatial clustering and long-range propagation. In contrast, SEP = 500 produces more localized and strongly attenuated responses, with smaller peak changes and limited spatial spread. These results support the conclusion that short excitatory projections substantially enhance network-wide propagation of single-neuron perturbations.

### Relationship between burst statistics and firing rate changes

To quantify how burst dynamics relate to perturbation-evoked responses, we computed, for each neuron, (i) the total number of detected bursts and (ii) the mean duration of its bursts across the simulation, and correlated these measures with the change in firing rate induced by single-neuron perturbation. Bursts were defined as sequences of at least three spikes with inter-spike intervals ≤ 7.5 ms, and correlations were assessed using Pearson’s product-moment coefficient across all neurons (for burst counts) or across all bursts (for burst durations).

**S2.**
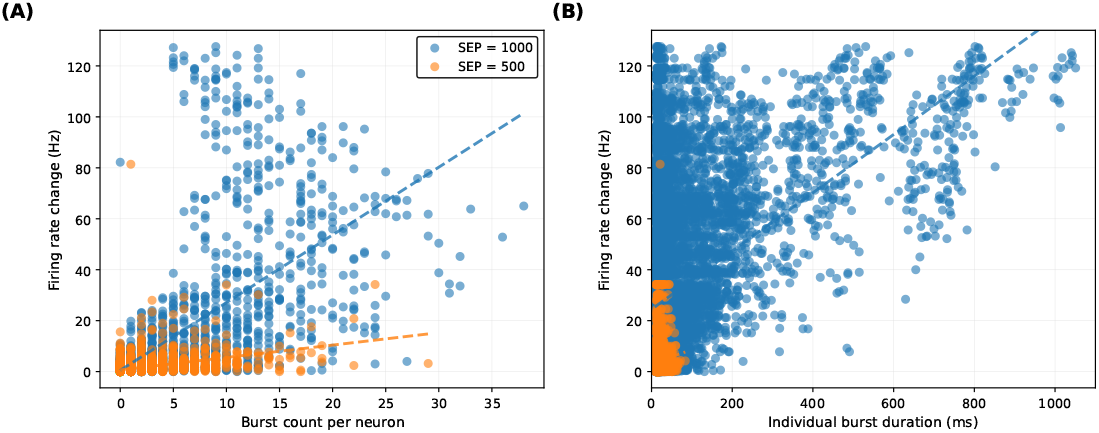
Burst activity and perturbation-evoked firing rate changes depend on short excitatory projections. Each point represents a single neuron (SEP = 1000, blue; SEP = 500, orange), and dashed lines indicate least-squares linear regression fits. **(A)** Burst count per neuron versus perturbation-evoked firing rate change. Networks with SEP = 1000 show a strong positive correlation (*r* = 0.677), whereas SEP = 500 exhibits a weaker but still positive correlation (*r* = 0.445), indicating that neurons with larger rate increases tend to generate more bursts, particularly in high-SEP networks. **(B)** Individual burst duration versus perturbation-evoked firing rate change. SEP = 1000 displays a moderate positive correlation (*r* = 0.469), while SEP = 500 shows only a very weak correlation (*r* = 0.084), suggesting that increased short excitatory projections strengthen the link between prolonged bursting and rate amplification. Bursts were detected using an inter-spike-interval threshold ≤ 7.5 ms and a minimum of three spikes per burst, and correlations were quantified using Pearson’s product-moment statistic.

### Structural modularity induced by short excitatory projections

To assess whether short excitatory projections (SEPs) induce structural modularity beyond their primary effect on functional organization, we computed Louvain modularity *Q* on the structural connectivity matrices of our networks. Networks were compared against a benchmark with 10 explicitly predefined modules (equal-sized, complete intra-module connectivity, no inter-module connections). Modularity was calculated across 100 independent network realizations per condition.

**S3.**
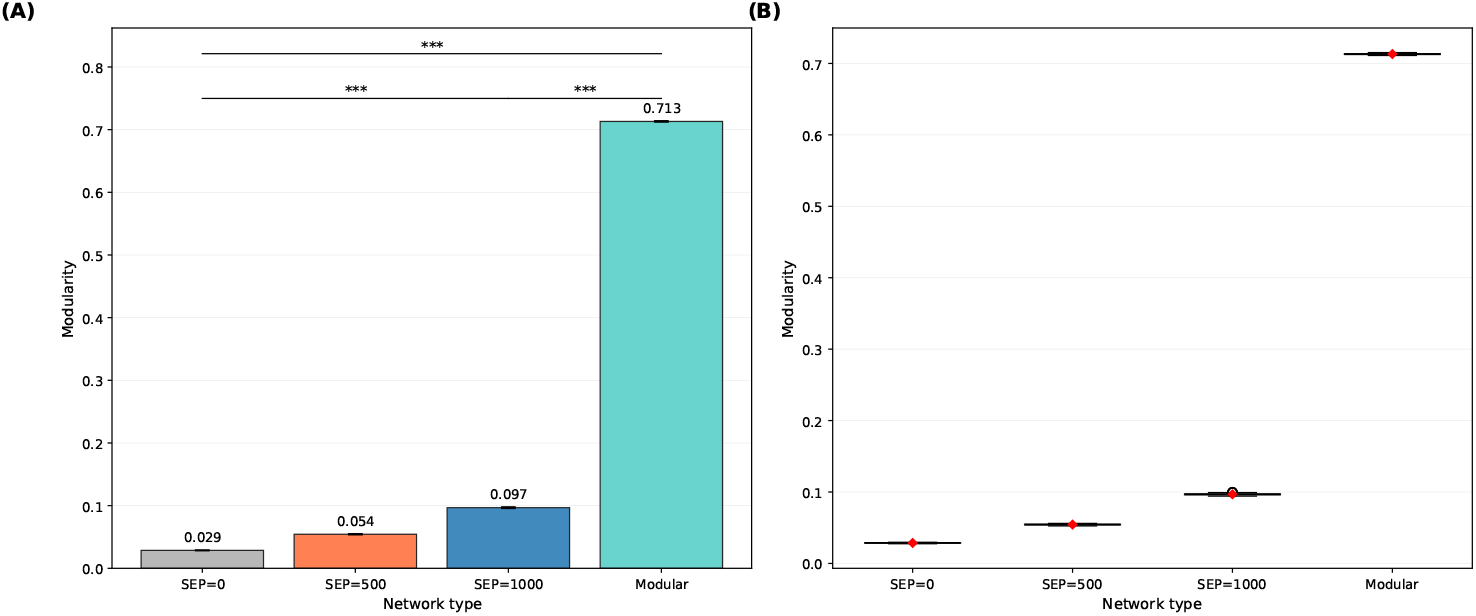
Short excitatory projections induce only weak structural modularity. Louvain modularity values for networks with different densities of short excitatory projections (SEP=0, SEP=500, SEP=1000) and for an explicitly modular benchmark network with 10 predefined modules. Bars (left) and boxplots (right) show modularity across 100 independently generated networks per condition (mean *±* SD). Modularity increases monotonically with SEP density (SEP=0 ≈ 0.03, SEP=500 ≈ 0.05, SEP=1000 ≈ 0.10), indicating the emergence of weak structural communities from an initially random baseline, whereas the benchmark modular network exhibits much higher modularity (≈ 0.71).

## Notes

### Competing Interest Statement

The authors have declared no competing interest.

